# Formalizing biodiversity measures and guiding through them

**DOI:** 10.1101/2025.08.25.672117

**Authors:** Sandrine Pavoine

## Abstract

Facing the myriad of indices claimed to be biodiversity indices that have been proposed by the scientific community, it becomes critical to analyze the mathematical properties that determine whether any index respond to biological necessities about biodiversity quantification. In ecology, species richness, the number of species, was a pioneer simple formula still currently most widely used as a biodiversity index. Species, functional, phylogenetic diversity indices then emerged from the feeling that combining species richness with the distribution of species’ abundances, functional and/or phylogenetic characteristics could provide indices that are more linked to ecological phenomena than species richness alone. Although there are theoretically an infinite number of potential biodiversity indices, I show here that some widely used indices of biodiversity actually take on large values in the absence of diversity. I thus claim for stricter definition of mathematical formulas that can be classified as biodiversity indices and for the use of parametric guiding indices that generalize and connect existing indices, tracing paths through the jungle of indices. As a case study I develop a parametric guiding index, and applying it on bat diversity, I show that analyzing the mathematical properties of potential biodiversity indices is critical to evaluate their relevance as regards a study objective in ecology. I come to the conclusions that to improve applications of biodiversity indices in ecology we need to: stop claiming for a single best formula; acknowledge that biodiversity indices are above all diversity indices, with the particularity that they are applied to biological data; agree on a set of minimal conditions that any biodiversity index must fulfill; integrate a large panel of diversity indices in software packages to enable ecologists to select and apply the most relevant index for their study. I propose a set of minimal conditions as a first step in this direction.

## Introduction

### On the importance of mastering the properties of biodiversity indices

The expression “biodiversity” is a general concept introduced in a political context linked to global changes during the National Forum of biodiversity that took place in 1986 (Wilson 1988). However the mathematical measurement of facets of biological diversity started long before 1986, so that many pivotal papers on the measurement of biodiversity date back from the 1960s and 1970s (e.g., MacArthur and MacArthur 1961, MacArthur 1965, Pielou 1966, McIntosh 1967, Hurlbert 1971, Whittaker 1972, Hill 1973, Peet 1974, Grassle et al. 1979). Here I simply define biodiversity in the literal sense as the variety of life. Measuring entire biodiversity with a single mathematical formula is an unachievable goal given the multiple aspects of diversity in life forms. This goes some way to explaining why scientific literature contains a multitude of mathematical formulae aimed to measure biodiversity. At this stage, I will designate any of these mathematical formulae as a potential biodiversity index. A basic requirement for any biodiversity index is that it is justified in both a biological and a mathematical context. The development of biodiversity indices must therefore be controlled by their properties, any property being “a biological or a mathematical necessity” (Hurlbert 1971). Applications of biodiversity indices in the ecological literature show that although the biological motivation is intuitively accepted by many, that of the mathematical necessity is often perceived as secondary. Yet, it is actually the mathematical properties that allow to check whether the biological motivations have been respected in the index formulation. Indeed, understanding the mathematical properties of biodiversity indices is an essential part of ecological research because these properties determine the way the index can be interpreted, the degree to which it responds to a biological necessity, and ultimately the finding of connections between biodiversity and ecosystem functioning, biodiversity and stability, biodiversity and resilience, etc. (e.g., Petchey et al. 2004, Ricotta 2006). The biological and mathematical necessities are thus intrinsically linked and may depend on the purpose of a study.

### The emergence of species diversity indices

In ecology, an intuitive way of quantifying biodiversity has early been to cluster organisms in species and count the resulting number of species (e.g., Agassiz 1854, DeCandolle 1855, Tristram 1897, Jaccard 1912). Hereafter I will use the general word “collection” to designate any set of species (e.g., within a community, a plot, a region). Species richness, the number of species in a collection, has been by far the most frequently used biodiversity index in ecology, and it is still dominantly used to study current biodiversity trends in a context of global change (e.g., Hillebrand et al. 2018). Yet, as soon as the 1950s, it was repeatedly claimed that species diversity increases not only with species richness but also with species evenness, i.e. the evenness in their abundance (which generally corresponds to the evenness in the number of individuals per species in a collection, even if other measures of abundance have been considered; Black et al. 1950, Odum et al. 1960, Lloyd and Ghelardi 1964, Pielou 1966, Hurlbert 1971, Hill 1973, DeBenedictis 1973). In 1971, Hurlbert considered the following reasons that have motivated the inclusion of abundance in species diversity indices: “(i) the observation that two collections could contain the same number of species and the same number of individuals but still have very different structures, and (ii) the intuitive feeling that the number of species and their abundances somehow could be combined into an index that would show a closer relation to other properties of the community and environment than would number of species alone.”

From this time onwards, the expression “species diversity indices” was used to designate mathematical formulas that depend only to, and increase with, the number of species and species abundance evenness. Many possible properties that a species diversity index could fulfill have been listed in the ecological literature. One of them, widely accepted, is sample size insensitivity: a species diversity index should not be functionally related to the number of individuals in a collection (Hubálek 2000). Indeed, the absolute abundance of all species does not count in species diversity indices, but relative abundances do. This is because individuals of the same species are considered identical. I focus on a list of four fundamental properties that concern the range of possible values for any index of species diversity (its minimum and its maximum possible value) and its sensitivity to species richness:

#### p0. Non-negativity

A species diversity index has non-negative values (e.g., Hubálek 2000).

#### p1. Full concentration at minimum

The species diversity of a collection reaches its minimum value when all individuals belong to (are concentrated in) one species (e.g., Hubálek, 2000), or more generally when species abundances are set to zero for all species, except one.

#### p2. Abundance evenness at maximum

Given a collection of species, the species diversity of the collection reaches its maximum value when its species are equally abundant (e.g., Margalef 1958, Patten 1962, Pielou 1966, Hurlbert 1971).

#### p3. Embedding of species richness

Given two collections in which the species are equally abundant, the collection with the largest number of species must have the highest value of species diversity (Hubálek 2000).

### From species diversity to functional and phylogenetic diversities

As soon as in the 1970s and more markedly in the 1980s and 1990s, while many researchers still promoted the consideration of species abundance in addition to the number of species in the measurement of biological diversity, others less numerous advocated the addition of species’ biological and ecological characteristics (e.g., morphology, niche) to biodiversity measures (e.g., Hendrickson and Ehrlich 1971, Rao 1982, Margalef and Gutierrez 1983, Vane-wright et al. 1991, Cousins 1991). The development of diversity indices that accounted for species’ biological and ecological characteristics, in addition to their abundance and number, has then gradually increased (see, e.g., Mouchet 2010, Tucker et al. 2017, Schmera et al. 2023, for reviews). Two aspects to characterize species have come to the fore in the ecological literature: the functional traits (traits that reflect a species’ function in its ecosystem or a species’ response to environmental changes, Lavorel and Garnier 2002), and the phylogenetic links (amount of evolutionary history shared by species, Nee and May 1997). When considering species’ biological characteristics, in addition to their abundance, whether phylogenetic links or functional traits, the development of biodiversity indices becomes more complex than that of species diversity indices. One can no longer rely on the same properties as for species diversity. For example, although the basic p0 property still seems reasonable, the p2 property cannot be characteristic of phylogenetic and functional diversity indices, as some species will be considered more similar than others.

Currently, few papers study the properties of functional and phylogenetic diversity indices. There is no rigorous framework for this development of indices. This has left the way open for the development and widespread use of indices with disregarded properties that are not always relevant for use in ecology. For example, an index widely used to measure functional diversity (FDiv, Villéger et al. 2008) can approach its maximum when a single species dominates in abundance (Kondratyeva et al. 2019). Knowing that individuals within species are assumed identical, it seems reasonable to consider conversely that a collection dominated by a single species has close to zero diversity (see also Kosman et al. 2021 and Supplementary material for another example).

### Objective

To guide the use of proposed functional and phylogenetic (hereafter FP) diversity indices, it becomes therefore necessary to (1) connect existing indices; (2) inform on their mathematical properties; (3) account for the context and objective of a study to specify expected biological necessities. To take a step in this direction, here I introduce a parametric index of FP diversity that connects and generalizes both proposed indices of FP diversity, that are Walker et al. (1999) index of functional attribute diversity (FAD), Schmera et al. (2009) modified functional attribute diversity (MFAD) index, Pavoine et al. (2017) R index, and Rao (1982) quadratic entropy index for FP diversity, and indices of species diversity including Gini-Simpson index (Gini 1912, Simpson 1949) and Guiasu and Guiasu (2010) Rich-Gini-Simpson quadratic index of biodiversity (Fig. 1). I propose to name a formula that links existing indices in the literature a “guiding parametric index”. A typical example of a guiding parametric index is Hill (1973) index that links species richness with Gini-Simpson and Shannon indices of species diversity.

**Figure 1.**
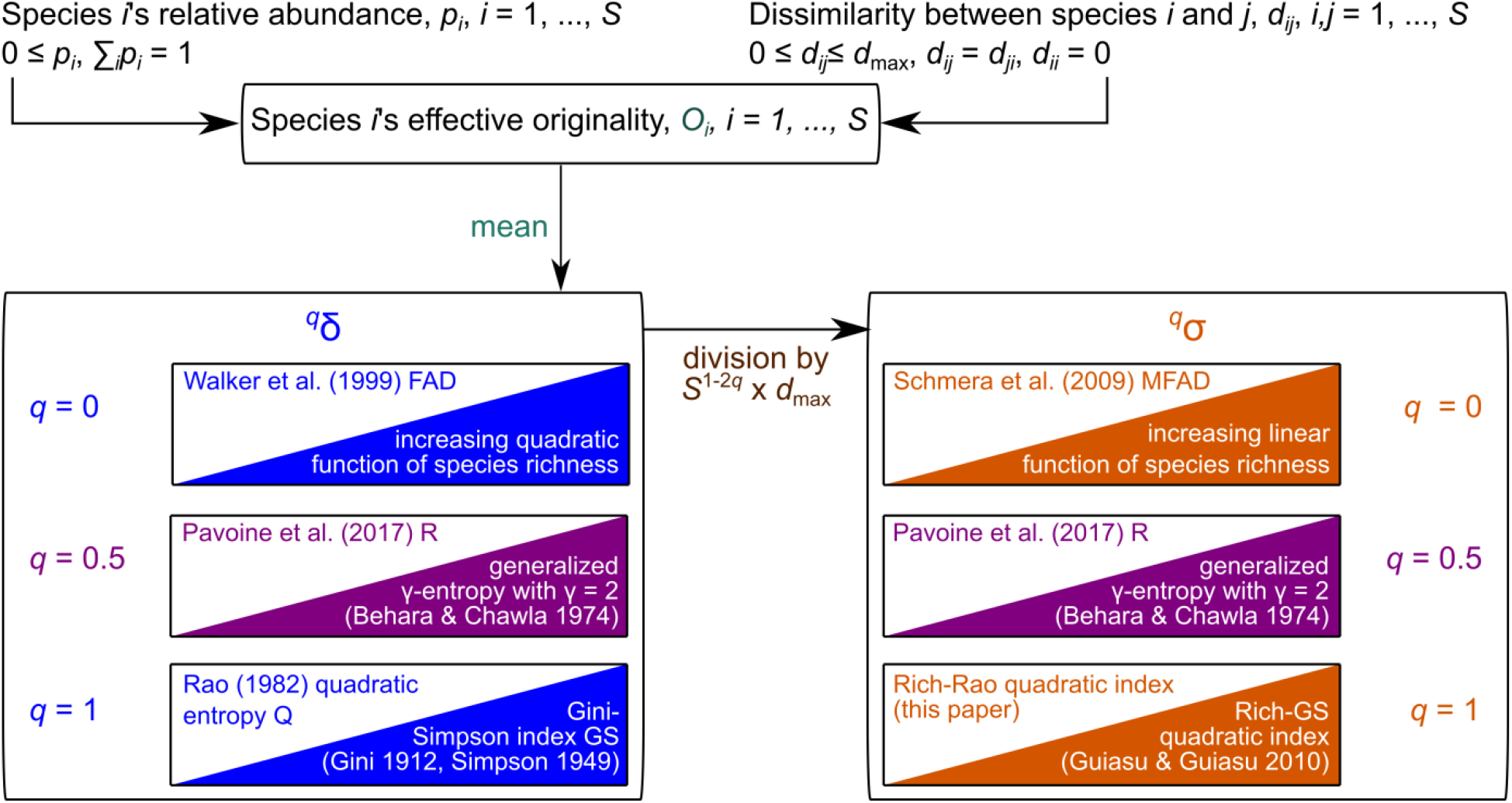
Guide through indices. The effective originality of a species depends on its dissimilarities with other species and on species’ relative abundance. ^*q*^*δ* is obtained as a mean of species’ effective originalities and ^*q*^*σ* as a standardization of ^*q*^*δ*. Particular cases of ^*q*^*δ* and ^*q*^*σ* are specified for *q* = 0, 0.5 and 1 as they relate to already published indices. Each link with existing indices is specified thanks to a rectangle containing: above diagonal, the related FP diversity index (for raw FP dissimilarities); and below diagonal, the related species diversity index (for equidissimilar species with *d*_*ij*_ = *d*_max_ for all *i* ≠ *j*).

## Methods

### Targeted properties

Noting that “Despite extensive literature on diversity-related issues, a formal definition and logical development of diversity as a concept and of its measurement have been lacking”, Patil and Taillie (1982) formalized the measurement of diversity by positing that diversity is an average property of a collection. At the time of their paper they identified that property as species rarity (a decreasing function of species abundance). Considering FP diversity, I identify that property as species effective originality. The effective originality of a species is the rarity, not of the species itself but, of its FP characteristics (be they rare functional traits or an isolated phylogenetic position, Pavoine and Ricotta 2021).

Here I consider that one can define a measure of (dis)similarity between two species by comparing any of their biological characteristics. The measure can lead to functional (dis)similarities (e.g., by using Mahalanobis distance on a quantitative table of species’ traits), or phylogenetic (dis)similarities (e.g., by calculating the age of the most recent ancestor common to two species). For the purpose of measuring FP diversity, I will name these (dis)similarities “FP (dis)similarities”. Here I show that the guiding parametric index of FP diversity I propose is a mean of effective originalities that satisfies the following properties:

#### P0. Non-negativity

The index has non-negative values.

#### P1. Full similarity at minimum

The index reaches its minimum value when species abundances are as unequal as possible (i.e., all individuals belong to one species) and/or when species are FP similar.

#### P2. Originality evenness at maximum

Given a collection of species with fixed FP dissimilarities between them, the index reaches its maximum value for the collection when its species have equal effective originality.

#### P3. Embedding of species diversity

When species are all FP equidissimilar, the index increases with this FP dissimilarity and satisfies the same properties as species diversity indices.

The multiplication of biodiversity indices evoked above also has its origins in the multiplicity of objectives for which we measure biodiversity. When this objective is preservation of biodiversity, we are influenced by what we want to preserve. Since the species is the basic unit in conservation discourse, a widespread thought is that we want to conserve as many species as possible (e.g., Parr et al. 2009, Wilson et al. 2011, Raven 2021, Carné and Vieites 2024). In this conservation biology context, “when each species counts”, I consider that one would expect that any index of species diversity and any index of FP diversity also fulfil the following properties respectively:

#### p4 (resp. P4). Weak species monotonicity

sensu Weikard et al. (2006). The loss of any species is accompanied by a loss of species diversity (resp. FP diversity) in comparison to the situation where the species is maintained with rare abundance.

I will therefore also study property P4 and identify when the guiding parametric index of FP diversity fulfills it.

### The guiding parametric index

Consider a collection of *S* species. Let **D** = (*d*_*ij*_)_1≤*i,j*≤*S*_ be the matrix of pairwise FP dissimilarities between species, with 0 ≤ *d*_*ij*_, *d*_*ij*_ = *d*_*ji*_ for all *i, j*, and *d*_*ii*_ = 0 for all *i*. Let **p** = (*p*_1_, …, *p*_*S*_)^t^ be a vector of species’ relative abundances (*t* being the transpose),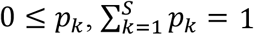. A guiding parametric index of diversity can be defined as

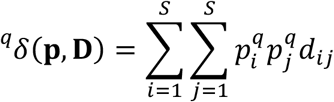

with *q* in [0,1]. This index ^*q*^ *δ* integrates the fact that an assemblage with a single species has zero diversity (given that within-species diversity is discarded). 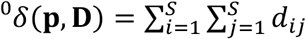 is related to Walker et al. (1999) index of functional attribute diversity (FAD).

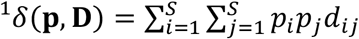 is Rao (1982) quadratic entropy (Q), the abundance-weighted average dissimilarity between two species in a collection. Index 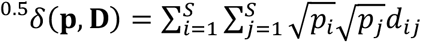 is Pavoine et al. (2017) R index. Thanks to Eq. 1, R can be here presented as an intermediate viewpoint between FAD and Q (Fig. 1). ^*q*^ *δ* thus combines Walker et al., Pavoine et al. and Rao indices and generalizes them. ^*q*^ *δ* is a mean of species effective originality: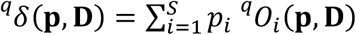, where

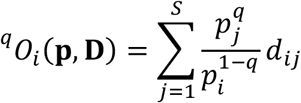

is the effective originality of species *i*. By its definition, ^*q*^*O*_*i*_ decreases with the relative abundance of *i, p*_*i*_; so it increases with its rarity. It also increases with the amount of difference between species *i* and other species, especially the most abundant ones. If species all are equidissimilar with *d*_*ij*_ = 1 for all *i* ≠ *j*, ^*q*^ *δ* reduces to an index of species diversity that satisfies properties p0 to p3 for all *q* in]0,1] and p4 for all *q* in]0, 1[(proof in Supplementary material)

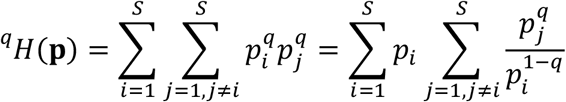

and the effective originality of any species *i* reduces to an index of relative rarity

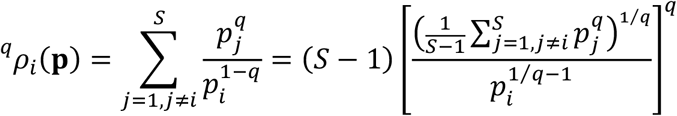

^0^*H*(**p**) = *S*(*S* − 1), is a function of species richness and 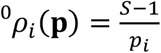 increases with species *i* rarity and with the number of other species in the collection. With index ^0^*ρ*_*i*_(**p**) a species is thus effectively original of it is rare among many other species. 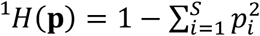 is Gini-Simpson index of species diversity and ^1^*ρ*_*i*_(**p**) = 1 − *p*_*i*_ is an index of rarity as proposed by Patil and Taillie (1982). With Gini-Simpson index, a species is thus considered effectively original provided it is rare. When 0 < *q* < 1, ^*q*^*ρ*_*i*_ increases with species *i*’s rarity, with the number of other species in the collection and the generalized mean (with exponent *q*) of the abundance of these species. When *q* = 1 the number of species does not count to define the effective originality of a species, while for *q* = 0 it is critical in this definition. Decreasing *q* from 1 to 0 is thus giving more and more importance to the number of species. As soon as *q* < 1, species richness counts as desired in a context of biological conservation. I will show in the next sections that this has important consequences on how to interpret and use the index ^*qδ*^.

### Weak species monotonicity

An index whose value never decreases when a species is added to a collection is said to be species monotonous (a property known as “species monotonicity” Weikard et al. 2006; or “set monotonicity” Izsák and Papp 2000). By definition any index of species richness must fulfill species monotonicity. However, unlike species richness, an index of species diversity that accounts for species abundances can decrease with the addition of a species in a collection as this addition may decrease abundance evenness. In this context, the property p4 defined above named “weak species monotonicity” (Weikard et al. 2006) ensures that, for any index of species diversity that fulfils it, there always exists a value of relative abundance sufficiently low but non-zero that ensures a species can be added to a collection without decreasing the value of the species diversity index.

When we account for the FP dissimilarities between species, if one add a species to a collection and this species is very similar to species already present, we can consider they increased diversity as they added a species, or we could consider that they decreased diversity by increasing the dominance of certain trait values or phylogenetic clades in the collection. Adapting the property of weak species monotonicity allows to reach a compromise between these two contrasting viewpoints: for any index of FP diversity that fulfils weak species monotonicity, there always exists a value of relative abundance sufficiently low but non-zero that ensures a species can be added to a collection without decreasing the value of the FP diversity index (property P4).

^*q*^ *δ* (**p, D**) fulfills species monitonicity when *q* = 0 and weak species monotonicity for all *q* in]0,1[(see proof in Supplementary material). It fails to satisfy this property when *q* = 1.

**Maximizing diversity…**

**…when only dissimilarities vary**

Maximizing ^*q*^ *δ* (**p, D**) over **D** (with the relative abundance vector **p** fixed) could be useful for*q* = 0, as by definition ^0^*δ* is insensitive to **p**, and for other values of *q*, when comparing communities that share the same abundance distribution but have different species with different FP characteristics. If the *d*_*ij*_ are not bounded then the maximum of ^*q*^*δ* (**p, D**) over **D** is ∞. In contrast, if, as in many ecological studies (e.g., Le Tortorec et al. 2023), the *d*_*ij*_ are bounded so that there exists a positive value *d*_*max*_ such that 0 ≤ *d*_*ij*_ ≤ *d*_*max*_, maximizing ^*q*^*δ* (**p, D**) over **D** is simply considering that the *d*_*ij*_ for all *i* ≠ *j* reach the maximum *d*_*max*_:

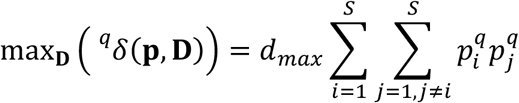

For *q* = 0, this yields

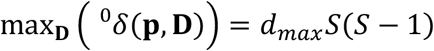

An index of FP diversity that varies between 0 and *S* - 1 and that does not account for species abundances thus is obtained by dividing ^0^ *δ* by *d*_*max*_ × *S*, leading to the standardized index:

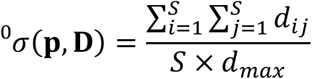

which linearly increases with Schmera et al. (2009) index 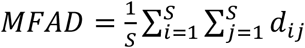 (Fig. 1). Note that *d*_*max*_ is often set to unity in ecology (e.g., Botta-Dukát 2005, Wayman et al. 2021, Le Tortorec et al. 2023) so that equations above and below should often simplify. Contrary to ^0^ *δ* (**p, D**), ^0^*σ*(**p, D**) does not fulfill species monotonicity as it can decrease by the addition of a new species to a collection (see Supplementary material for an example). When dissimilarities are even (*d*_*ij*_ = *d*_*kl*_ for all *i* ≠ *j, k* ≠ *l*), ^0^*δ* and ^0^*σ* reduce to a quadratic and to a linear function of species richness, respectively (Fig. 1).

**…when abundances and dissimilarities both vary**

Maximizing ^*q*^ *δ* (**p, D**) over **D** and **p** is useful for *q ≠* 0 when comparing communities, e.g.,from different regions, that have neither the same abundance distributions nor the same species. The maximum of ^*q*^*δ* when both **p** and **D** vary is obtained when the *S* species of the collection have equal relative abundance, i.e. *p*_*i*_ = 1/*S* for all *i*, and *d*_*ij*_ = *d*_*max*_ (the maximum possible dissimilarity between two species) as soon as *i* ≠ *j*. This maximum is:

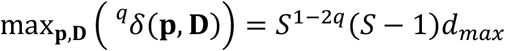

The sensitivity of ^*q*^*δ* to species richness thus depends on *q*. Only when *q* = 0.5, the maximum linearly increases with species richness.

When *q* ≠ 0, ^*q*^*δ* can be standardized as follows so that, when both abundance and dissimilarity vary, the resulting standardized function ^*q*^*σ* varies between 0 and *S* - 1, having thus a maximum that linearly increases with species richness:

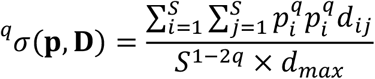

When *q* = 0.5,

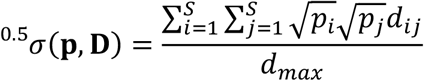

is proportional to ^0.5^ *δ*(**p, D**) (=Pavoine et al. 2017, R index). ^0.5^*δ* reduces to γ-entropy with γ = 2 (Behara and Chawla, 1974) if species are maximally dissimilar (Fig. 1).

When *q* = 1,

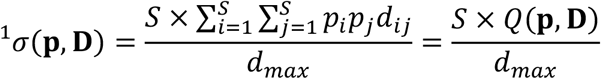

increases with Rao’s quadratic entropy and species richness, and reduces to Guiasu and Guiasu (2010) Rich-Gini-Simpson quadratic index of biodiversity if species are maximally dissimilar. A Rich-Rao quadratic index of biodiversity can thus be introduced here as a generalization of Guiasu and Guiasu (2010) Rich-Gini-Simpson quadratic index: *RQ*(**p, D**) = *S*× *Q*(**p, D**) (Fig. 1).

In a collection where species are maximally dissimilar, ^*q*^*σ*(**p, D**) reduces to

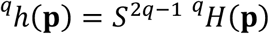

^*q*^*h, q* in]0,1], is an index of species diversity that satisfies properties p0 to p3 for all *q* in]0,1] and p4 for *q* in [0.5,1] (proof in Supplementary material).

**…when only abundances vary**

Maximizing ^*q*^ *δ* (**p, D**) over **p** is useful for example when one considers a fixed regional pool of species and wonders what would be the composition of a collection, drawn from this pool, that would be as diverse as possible. Dissimilarities between species in the species pool are fixed and cannot change within the collection, only abundance can vary. Maximizing ^*q*^*δ*(**p, D**) when only **p** varies also gives a better understanding of the behavior of the function ^*q*^*δ*. ^0^*δ* is insensitive to **p** and will thus not be studied here.

As far as *q* = 1, at the maximum of ^1^*δ*(**p, D**) over **p**, a species is either absent (null abundance) or is present (positive abundance) with an effective originality equal to that of the other present species. The property weak species monotonicity ensures that for *q* in]0,1[all species are present at the maximum. For all *q* in]0,1[, ^*q*^ *δ* (**p, D**) index reaches its maximum value over **p** when the species in the collection all have equal effective originalities (proof in Supplementary material).

In the previous section, I introduced a normalized version of ^*q*^*δ* (**p, D**), 0 < *q* ≤ 1, so one can legitimately wonder how the normalized version ^*q*^*σ*(**p, D**) behaves when only **p** varies, while **D** is fixed. For *q* = 0.5, ^0.5^*σ* = ^0.5^*δ*/*d*_*max*_, so that ^0.5^*σ* and ^0.5^*δ* have similar behavior. For 0.5 < *q* < 1, ^*q*^*σ* fulfills weak species monotonicity. The maximum of ^*q*^*σ* is the same as that of ^*q*^ *δ*. This is because ^*q*^*σ* is an increasing function of both species richness and ^*q*^*δ*, and the maximum of ^*q*^ *δ* is reached when species richness is also maximum. For *q* = 1, let π_*i*_ be the relative abundance of any species *i* at the maximum of ^1^ *δ* and *n* the number of species that have zero abundance at the maximum of ^1^*δ*. Then, ^1^*σ* tends to its maximum when the relative abundance of *i* is *ε* if *π*_*i*_ = 0, and *n*_*i*_ × (1 − *n* × *ε*) if π_*i*_ ≠ 0, with *ε* tending to 0^+. 1^*σ* and thus also the Rich-Rao quadratic index of biodiversity therefore tend to their maximum without excluding species, as all species have non-zero abundance. However, certain species are extremely rare. When 0 < *q* < 0.5, ^*q*^*σ* increases with ^*q*^*δ* but decreases with species richness. In contrast to ^*q*^*δ*, when 0 < *q* < 0.5, ^*q*^*σ* does not fulfill weak species monotonicity and there exist some matrices **D**, such that ^*q*^*σ*(**p, D**) is maximized over **p** for a vector of relative abundance that contains at least one zero (see an example in Supplementary material).

### Synthesis of satisfied properties

Properties P0 to P4 are generally satisfied by ^*q*^*δ* and ^*q*^*σ* except in the following cases: the elimination of a species from the collection may sometimes increase ^*q*^*σ* when 0 ≤ *q* < 0.5 and ^*q*^ *δ* when *q* = 1, yielding P4 unsatisfied and P2 only partially satisfied (Fig.2).

**Figure 2.**
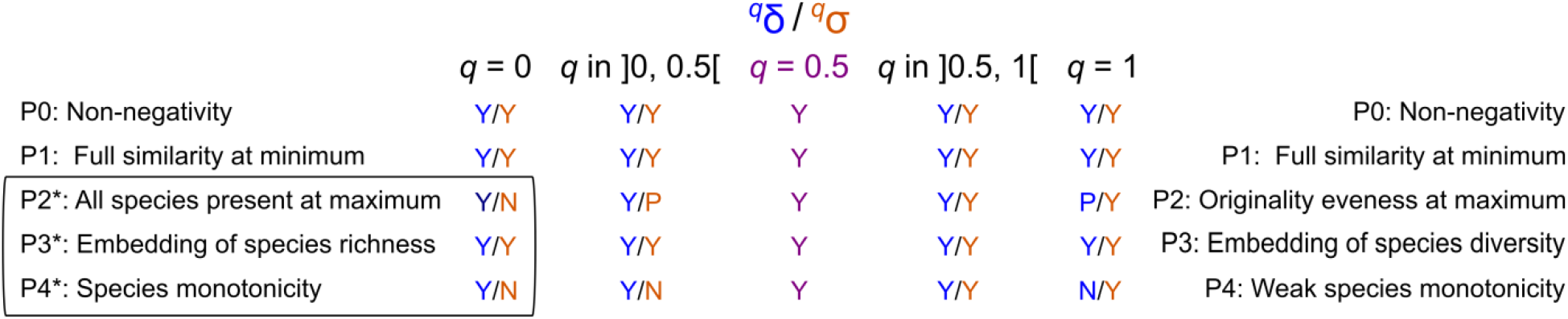
Synthesis of the properties satisfied by ^*q*^*δ* and ^*q*^*σ*. Properties P0 to P4 are those detailed in the Introduction. Properties P2 to P4 are only valid for indices which take abundances into account. Properties P2*, P3* and P4* are thus adaptations of the properties P2, P3 and P4, respectively, for an index that does not take species’ abundances into account (i.e., ^0^*δ* and ^0^*σ*). For P1, full similarity means maximal similarities between individuals for indices that consider abundances, and maximal similarities between species for indices that doesn’t. A property can be satisfied (Y for yes); not satified (N for no); or partially satisfied (P for partially). “Partially” is used for P2, when the index can be maximized for a subset of evenly original species with others having zero abundance. To be fully satisfied, P2 excludes null abundances at maximum. P2 and P2* considers maximization over all possible distributions of presence (for *q* = 0) or abundance (for 0 < *q* ≤ 1) of species, with fixed dissimilarities between species.

### Case study

#### Data

Medellín et al. (2000) observed and identified bat species in four habitats (rainforest, cacao plantation, old field, cornfield) in the Selva Lacandona of Chiapas, Mexico. Rainforest was dense with tall trees, lianas and understory. Cacao plantations were composed of cacao trees (*Theobroma cacao*) but also retained several tall trees of the rainforest as they provide shade to cacao trees. Old fields were abandoned agricultural fields covered with secondary vegetation. Cornfields were active corn-dominated fields characterized by a vegetation that was neither diverse nor structurally complex (see Medellín et al. 2000, for more details on the data set). To evaluate morphological differences between observed species, I considered three morphological traits: head-body length (distance from anus to muzzle tip), tail length (distance from anus to tail tip), ear length (distance from the notch to the fleshy tip of the pinna). Values were obtained from Wilson and Mittermeier (2019) thanks to platform Treatment Bank (plazi.org). When an interval of values was specified, I considered its center as reference value.

#### Analyses

I analyzed bat morphological diversity in these habitats for an even number of observed individuals. For that I retained all the 444 individuals observed in rainforest (smallest sample) and randomly sampled 444 individuals in each other habitat. I also randomly sampled 444 individuals over all habitats and 444 individuals over the agricultural habitats, to roughly evaluate the influence a mosaic of habitats could have on the overall diversity of a region. This approach of sub-sampling allows evaluating levels of diversity independently of the absolute abundance of bats. I then calculated the morphological distances between species with Mahalanobis distance applied to the three morphological traits to account for potential partial correlations between the traits.

I calculated the morphological diversity within each sample using ^*q*^ *δ* and ^*q*^*σ*, with *q* varying from 0 to 1 with a step of 0.1. For *q* varying from 0.1 to 1 with a step of 0.1, I also determined the theoretical vectors of species’ relative abundances that maximize ^*q*^ *δ* and ^*q*^*σ* applied to the morphological distances between species. Optimization was done with R package Rsolnp version 2.0.0, function csolnp (Galanos and Ye 2025, R Core Team 2025).

#### Results

The sample drawn from rainforest always appeared the most diverse and that sampled from cornfields the least diverse. The ranking of the other samples depended on the index used to measure diversity (Figs 3-4). For low values of *q* (*q* < 0.4111 [threshold are given here rounded with 4 digits]), using ^*q*^*δ* or ^*q*^*σ* led to the following ranking from most diverse to least diverse sample (for an equal number of observed individuals): any habitat > old fields > cacao plantation > any agricultural field. However this ranking changes when increasing *q*: for 0.4111 < *q*, the diversity in cacao plantation was perceived as higher than that of old fields; as soon as 0.6829 < *q*, that of the sample drawn over all agricultural fields also exceeded the diversity of old fields.

**Figure 3.**
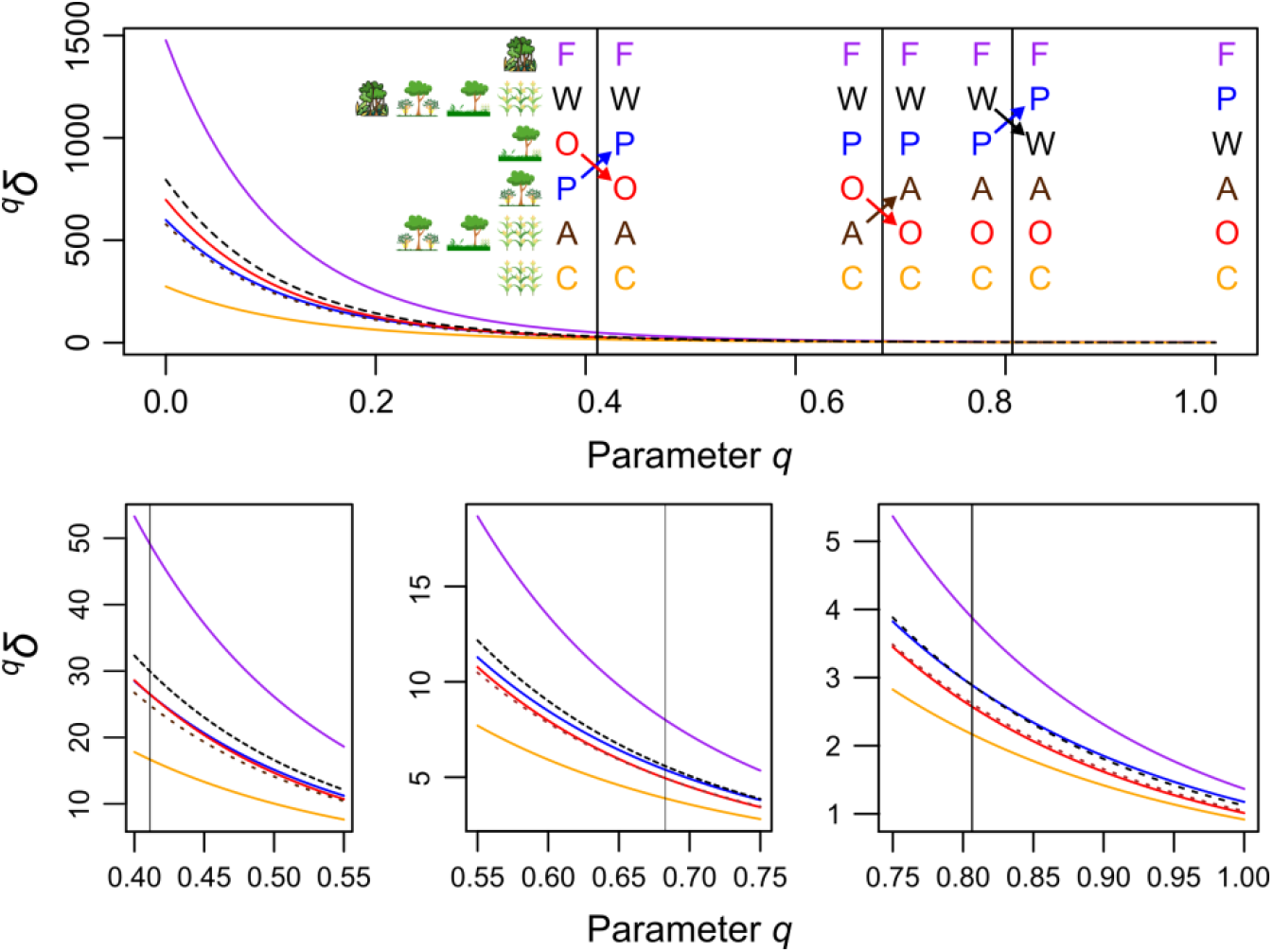
Diversity index ^*q*^*δ* applied to each sample of bats as a function of parameter *q*. Samples correspond to species observed in rainforest (F), in cacao plantations (P), in old fields (O), in cornfields (C), in any of the agricultural habitats (A = P, O, and/or C) and in any of the habitats (W = F, P, O, and/or C). The samples are sorted in the first panel from top to bottom from greater to lower diversity. Vertical lines indicate where the sorting of the habitat changes; and arrows indicate how it changes. Bottom panels are zooms around these changes.

The only difference in habitat ranking between ^*q*^*δ* or ^*q*^*σ* was obtained for high values of *q*: for *q* > 0.8065, using ^*q*^*δ* led to the sample obtained from cacao plantation being more diverse than the sample drawn over all habitats (Fig. 3); while for ^*q*^*σ* the sample obtained from cacao plantation never exceeded that drawn over all habitats (Fig. 4).

**Figure 4.**
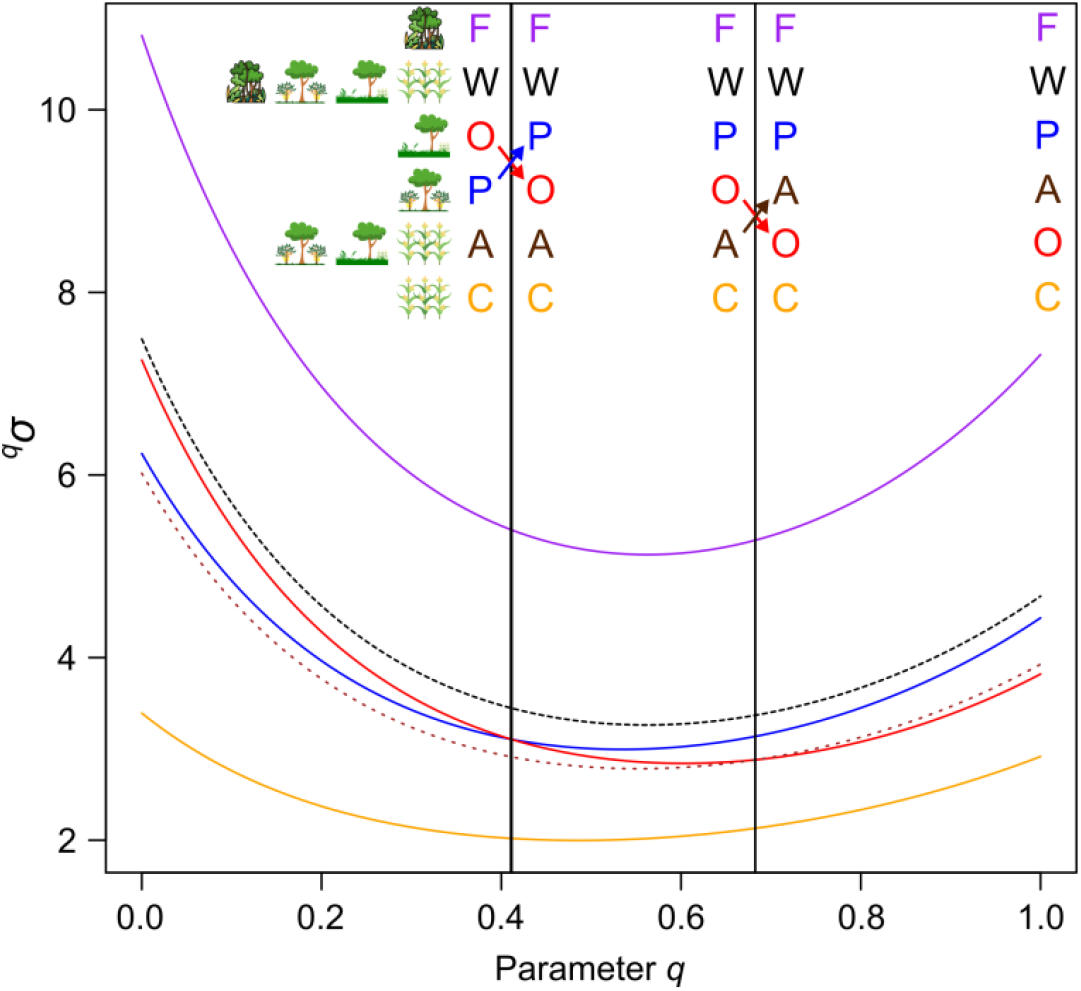
Standardized diversity ^*q*^*σ* applied to each sample of bats as a function of parameter *q*. Samples correspond to species observed in rainforest (F), in cacao plantations (P), in old fields (O), in cornfields (C), in any of the agricultural habitats (A = P, O, and/or C) and in any of the habitats (W = F, P, O, and/or C). The samples are sorted in the figure from top to bottom from greater to lower diversity. Vertical lines indicate where the sorting of the habitat changes; and arrows indicate how it changes.

For this case study, the vector of species’ relative abundance that maximizes ^*q*^*σ* was similar as the vector that maximizes ^*q*^*δ* for *q* = [0, 1[. For *q* = 1, as expected, let π_*i*_ be the relative abundance of any species *i* at the maximum of ^1^*δ* and *n* the number of species that have zero abundance at the maximum of ^1^ *δ*. Then, ^1^*σ* tends to its maximum when the relative abundance of *i* is *ε* if *π*_*i*_ = 0, and *n*_*i*_ × (1 − *n* × *ε*) if π_*i*_ ≠ 0, with *ε* tending to 0^+^. For *q* = 1, six species dominate in abundance at the maximum (Fig. 5). These are: Van Gelder’s bat *Bauerus dubiaquercus* characterized by the longest tail and long ears compared to head-body size; Big-eared woolly bat *Chrotopterus auritus* characterized by the largest head-body size and long ears; hairy-legged vampire bat *Diphylla ecaudata* characterized by large head-body length, small ears relative to head-body length and absence of tail; hairy-legged myotis *Myotis keaysi* characterized by an average head-body size, small mouse ear, and the second largest tail relative to head-body size; Davis’s round-eared bat *Tonatia evotis* characterized by an average head-body size, long tail, and the longest ears relative to head-body size; southern little yellow-eared bat *Vampyressa pusilla* characterized by its small head-body size, the absence of tail, and average ear relative to head-body size. As *q* decreases, *B. dubiaquercus, C. auritus, M. keaysi* and *T. evotis* (Fig. 6) remain the top abundant species at the maximum, all other species having nevertheless non-negligible abundance (Fig. 5). *D. ecaudata*, however, goes from the 4th highest to the 12th highest abundance at the maximum of ^*q*^*δ* and ^*q*^*σ*, when *q* decreases from 1 to 0.1; and *Vampyressa pusilla* from the 6th highest to the 22th highest abundance. Abundance of all species tend to become even at the maximum when *q* approaches 0 (Fig. 5).

**Figure 5.**
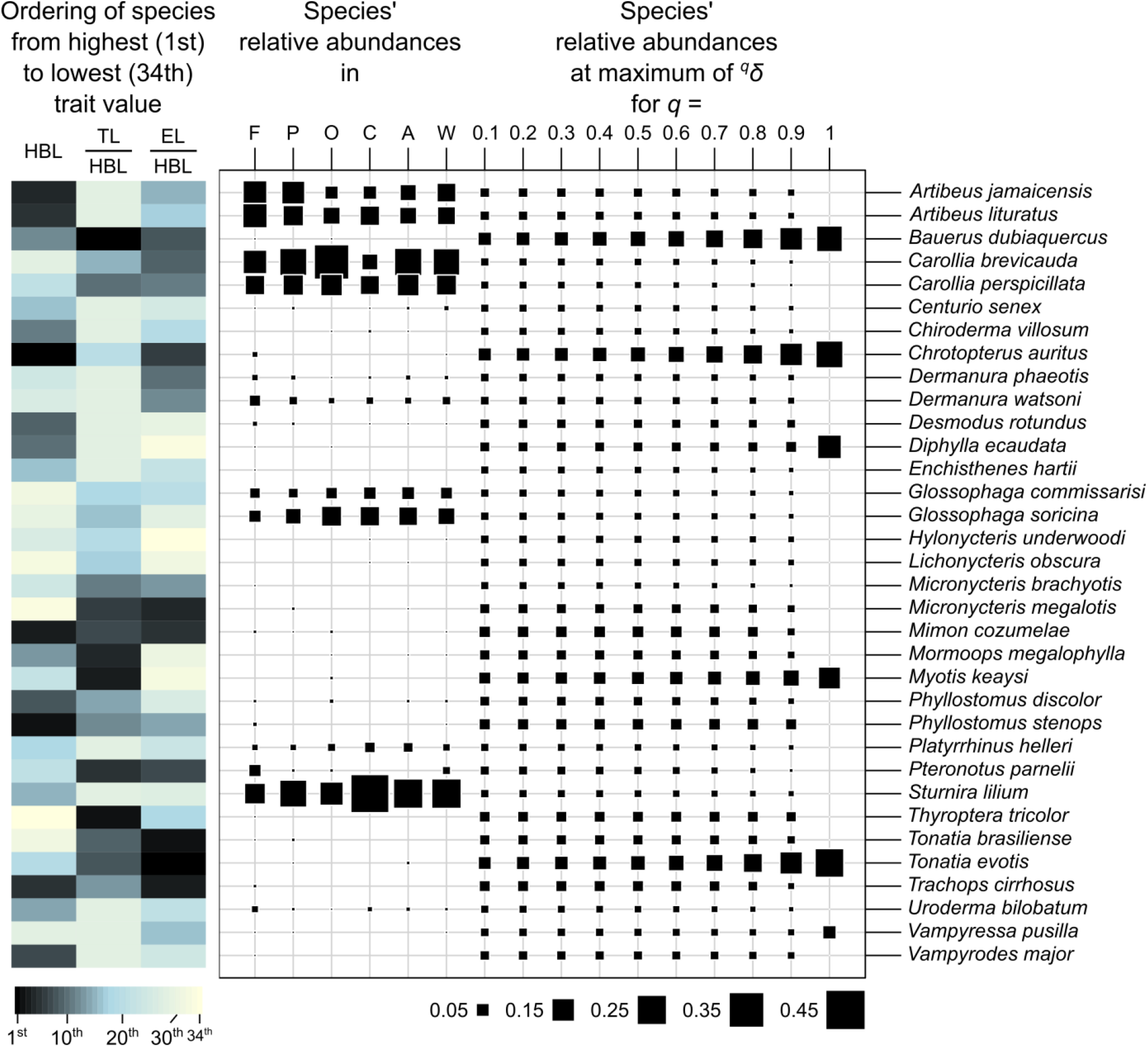
Abundance and traits of species in the case study. Left table: morphological description of the species with head-body length (HBL), ratio of tail length (TL) to HBL, ratio of ear length (EL) to HBL; color scale ranges from darkest (1st highest trait value) to lightest (34th rank; lowest trait value). Right table: Species relative abundance either within samples (F, P, O, C, A and W) or at the maximum of ^*q*^*δ*, with *q* ranging from 0.1 to 1 with a step of 0.1. Samples contain species from rainforest (F), cacao plantations (P), old fields (O), cornfields (C), any of the agricultural habitats (A = P, O, and/or C), and any of the habitats (W = F, P, O, and/or C). Species abundance is represented by the size of a black square from the absence of a square (species’ relative abundance = 0) to the largest square (relative abundance of *Sturnira lilium* in cornfields = 0.419).

**Figure 6.**
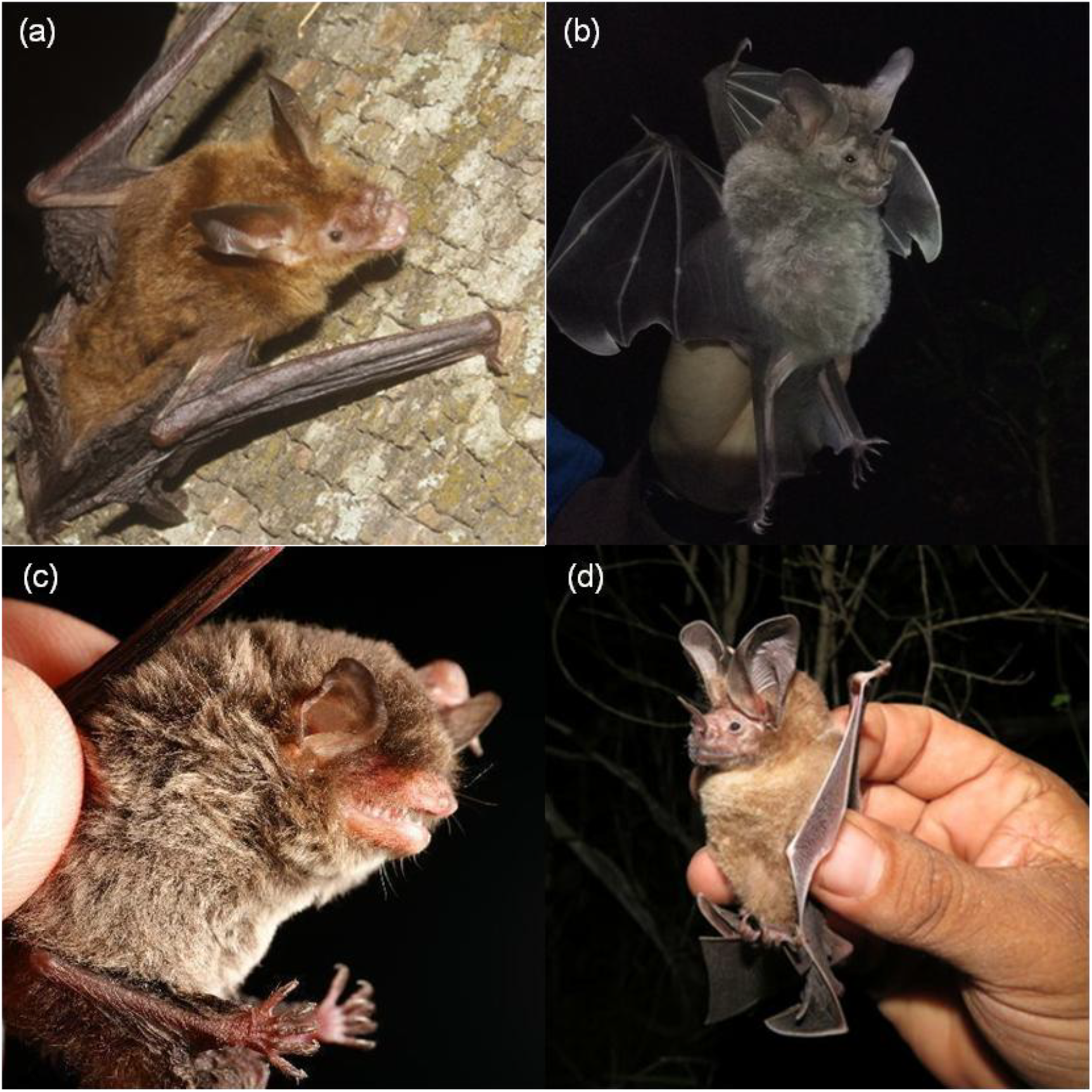
Photos of the four species that remain among the most abundant for maximizing morphological diversity in our case study (as measured with ^*q*^*δ* and ^*q*^*σ*, with *q* from 0.1 to 1 with a step of 0.1; see Fig. 5): (a) *Bauerus dubiaquercus*; (b) *Chrotopterus auritus*; (c) *Myotis keaysi*; (d) *Tonatia evotis*. Licences for photos (from top left to bottom right): CC BY-SA (author: Juan Cruzado Cortés); CC BY 4.0 (author: Nathan Van Cooten); CC BY-SA (author: Alex Borisenko); CC BY 4.0 (author: Giovana A. Valencia). The photos have not been edited other than adjusting their size (zoom on bats).

## Discussion

The approach above illustrates how the development of biodiversity indices has to be concomitant with the explanation and analysis of their properties. By developing ^*q*^*δ* and ^*q*^*σ*, I focused on properties P0 to P4. The properties of these indices could be studied in even greater depth. Indeed, different ecological conclusions were obtained depending on whether ^*q*^*δ* or ^*q*^*σ* was used to measure morphometric diversity and depending on the value of parameter *q* for these indices. Despite the clear cut results obtained for rainforest (perceived as the most diverse habitat) and cornfield (perceived as the least diverse habitat), uncertainty remained on the ranking of other habitats. This uncertainty can be removed and the different rankings explained by acknowledging the role *q* plays in ^*q*^*δ* and ^*q*^*σ* and the differences in ^*q*^*δ* and ^*q*^*σ* equations. For low (resp. high) values of *q*, ^*q*^*δ* and ^*q*^*σ* are more (resp. less) affected by changes in the abundance of rare species than by changes in the abundance of common species. This is illustrated by the fact that, in the case study, abundance evenness increased in the vector of species’ relative abundance that maximized ^*q*^*δ* and ^*q*^*σ* as *q* decreased. The high relative abundance of the non-distinctive silky short-tailed bat *Carollia brevicauda* in oldfields explain why for high values of *q*, oldfield was perceived less diverse than cacao plantation and than a mix of agricultural habitats. In addition, as *q* increases, species richness less and less contribute to ^*q*^*δ*. For increasing *q* > 0.5, species richness increasingly contribute to the value of ^*q*^*σ* in comparison to its impact on ^*q*^*δ*. This explains that for high values of *q*, ^*q*^*σ*, contrary to ^*q*^*δ*, did not classify the sample obtained from cacao plantation with 19 species as more diverse than the sample drawn over all habitats that totalized 21 species.

Research into the properties of diversity indices was very common on an empirical or analytical basis when it came to measure species diversity (e.g., DeBenedictis 1973, Heip and Engels 1974, Peet 1974, Kempton and Wedderburn 1978, Laxton 1978, Routledge 1979). While diversity measures have been largely extended to other (functional and phylogenetic) aspects of biodiversity, research into the properties of diversity indices are less frequent today. The development of biodiversity indices thus suffers from the lack of formal evaluation of newly developed indices before their widespread applications. This may be due to the lack of a reference set of properties that diversity indices should satisfy. The problem is that there is no limited list of properties that a biodiversity index must verify because of the multiplicity of purposes for which biodiversity indices are used. For example, property P4 studied in this article may be considered necessary in conservation biology, where the aim often is the preservation of the most diverse areas. There, each species counts: those involved in preserving biodiversity, and increasingly the general public, are generally concerned by the premature loss of a species, especially if it is driven by human activities. On the other hand, property P4 may be considered undesirable in community ecology, where biodiversity indices are used to understand how communities assemble. Indeed, this P4 property leads to indices that are highly correlated with species richness. The redundancy between species richness and the FP indices becomes a constraint to be managed for, and even an impediment to, the detection of certain ecological mechanisms (e.g., Sandel 2018).

Until the 1980s, the term biodiversity was not yet fashionable. It was simply referred to as diversity. The question of limiting the definition of this word ‘diversity’ to make it useful for ecological scientific research was already being raised (e.g., Hurlbert 1971, Peet 1974). Now that the word biodiversity is largely used both in the media and within scientific communities, the development of biodiversity indices suffers from the lack of formal rigor in the use of the term biodiversity. Those days, the term biodiversity is frequently used non-technically as a synonym for life, nature or wilderness, both in the non-scientific media and in the scientific community (Contoli and Luiselli, 2015). These shortcuts that equate biodiversity with ‘bio’ and omit the critical ‘diversity’ (the variable that should be targeted) opens the way to any mathematical formula applied to biological data to be qualified as a biodiversity index. This trend is even more pronounced when these formulae are developed with the aim to be used as biodiversity status indicators for public policy; there, a great deal of effort is being made to put “life in equation” without necessarily focusing on diversity. On the contrary, a biodiversity index must be first and foremost a diversity index, with the particularity that it is applied to biological data. ‘Diversity’ is the variable to be measured and ‘bio’ the data source set on which to measure it.

The development of biodiversity indices has suffered from the fact that many ecologists asked for a unique mathematical formula that could be used to measure diversity in an ecological context: e.g., “Despite the semantic and conceptual problems, a single summary statistic that will suitably characterize the species diversity in a sample (area, habitat, community, etc.) continues to be highly appealing to most ecologists because it makes a simple comparison of samples (communities etc.) possible.” (Hubálek 2000). If ‘biodiversity’ has to be something we can measure by a single mathematical formula representing all aspects of life variety, then biodiversity does not exist: “Communities having different species compositions are not intrinsically arrangeable in linear order on a diversity scale. Diversity per se does not exist.” (Hurlbert 1971). For biodiversity to be a meaningful and useful concept, we have, in contrast, to let it be multidimensional, acknowledging the impossibility to summarize it by a single quantitative score.

The development of multiple biodiversity indices is however perceived as an annoyance by the ecological community struggling to find its way through a jungle of indices to the most appropriate index for a given study (Ricotta 2005). The multiplicity of indices and the lack of explanation of their properties have indeed made it difficult to select an index for a given study. Hurlbert (1971) early identified that diversity indices used in ecology were those that were promoted by scientific figures and that have become fashionable. More recently, an additional reason explains the preferred use of certain indices: the ease of calculation in a free, easy-to-use software package. The most widely used indices are therefore the easiest to find (because they are promoted, fashionable) and to calculate (because the software is available).

This leads to the following conclusions: **First, biodiversity cannot be measured by a single formula: we need several biodiversity indices; Second, biodiversity indices are diversity indices applied to biological data; Third, to be clear on what “biodiversity indices” designate, we need a set of minimal conditions that any biodiversity index must fulfill; Four, we need index-rich software packages to enable ecologists to select and apply the most relevant index for their study**. In the last part of the discussion below, taking the case study I developed in this paper as a reference, I propose to tackle the first three points of this conclusion on which any software development rely.

From a mathematical viewpoint, biodiversity has thus to be viewed as a variable, the diversity, apprehended on a complex set of entities, the living. The living is a set that can be partitioned in different ways into smaller elements (building blocks). Each part can be subject to a measure of diversity. It remains to be clarified what is meant by diversity. Diversity is the character of that which is varied and thus of that which contains differences. Indeed, we cannot talk about diversity without talking about differences: diversity results from the existence of differences. To simplify the writing of minimal conditions, I thus consider that diversity indices are applied to sets of basic entities between which we can measure differences and conversely similarities. I thus propose the following minimal conditions that any diversity index applied to a set of entities should satisfy:

C1. A diversity index is nonnegative.

C2. A diversity index decreases with the similarity between entities.

C3. A diversity index amounts 0 when all entities are identical or when the set contains a single entity.

C4. If *d*_*ij*_ quantifies the differences between any 2 entities *i* and *j*, a diversity index reduces to a monotonically increasing function of the number of entities if these are equidifferent (i.e., *d*_*ij*_= *d*_*kl*_, for all *i* ≠ *j, k* ≠ *l*).

These conditions imply that if there are groups of identical species, then the number of groups is important in diversity measurement in addition to the relative abundance of each group. Indeed, if there exist groups of identical entities, then the imbalance of abundance between groups yields the increase of the overall perceived similarity between entities. This is typically the case in ecology when individuals are grouped by species and individuals of the same species are considered identical.

These conditions also imply that, if the basic entities are species, species are equidifferent and *N* is the number of species in a collection, then *H* = *N* - 1 can be considered as an index of biodiversity while *R* = *N* itself cannot. Indeed if *N* = 1, then *R* = 1, whereas there is no diversity in a set with a single entity; *H* instead correctly takes value 0.

Any biodiversity index applied to a set of entities should satisfy conditions C1-C4 above in addition to:

C5. A biodiversity index is measured on biological data collected from biological entities.

In this paper I considered typical biodiversity indices used in ecology where entities are individual organisms grouped by species. These indices depend the vector **p** of relative abundances, describing how organisms are distributed between species and the matrix **D** that provides the degree of dissimilarity between organisms from different species. Organisms from the same species are considered identical. The guiding parametric indices ^*q*^*δ* and ^*q*^*σ* take non-negative values. ^*q*^*δ* and ^*q*^*σ* always decrease by decreasing values in **D** (at least one value) until reaching 0 if **D** contains only zeros. By increasing the number of individuals of the already most abundant species, ^*q*^*δ* and ^*q*^*σ* decrease (because the number of identical organisms increases in the collection), until tending towards 0 when the relative abundance of the species tends towards 1. If each species was represented by a single organism and if species were equidissimilar, then both ^*q*^*δ* and ^*q*^*σ* would be increasing functions of the number of species. ^*q*^*δ* and ^*q*^*σ* thus fulfill all conditions C1-C5 above.

The definition of minimal conditions is therefore important to formalize the use of the expressions ‘diversity index’ and ‘biodiversity index’. The verification of these minimal conditions should not, however, prevent further study of the properties of each proposed (bio)diversity index. For example in our case study, the ranking of habitats according to their diversity depended on whether index ^*q*^*δ* or ^*q*^*σ* was used and on the choice for the value of *q*. Indeed, ^*q*^*δ* or ^*q*^*σ* have different mathematical properties I explicated and varying *q* also modifies these properties. Without studying the behavior of diversity indices through their mathematical properties, the conclusions of studies based on the interpretation of these indices’ values may be misleading and become meaningless. It is also necessary to continue to connect the indices already proposed as done here through parametric guiding indices to guide users within this perceived jungle of (bio)diversity indices.

## Scripts and data are available from

Figshare: https://doi.org/10.6084/m9.figshare.29979859.v1

