## Supplementary material for "Formalizing biodiversity measures and guiding through them"

### No diversity without differences

Diversity is "the fact of many different types of things or people being included in something" (dictionary.cambridge.org/dictionary/english/diversity, accessed August 12, 2025).

**Some indices proposed in the literature as diversity indices cannot be qualified as such because they approach their maximum value when almost all the entities in a collection are identical. By that, they do not respect the condition C2 specified in the Discussion section of the main text: A diversity index decreases with the similarity between entities.**

On a more general note, these indices fail to adhere the definition of the word 'diversity'.

An example is given in the main text. Another here:

The abundance weighted MNTD $\sum_{k=1}^{S} p_k d_{k\mathrm{min}}$ (Webb et al. 2002, Kembel et al. 2010) is maximized by retaining only the 2 most different species (whatever their abundance). It is thus not an index of diversity as it can be close to maximum when a single species dominates in abundance.

### Proofs

#### FAD increases by mixing

Let Ω be a reference set of species.

Let A be the set of species in a community a, $\overline{\mathrm{A}}$ be the set of species in Ω that are not in A; similarly, let B be the set of species in a community b, $\overline{\mathrm{B}}$ be the set of species in Ω that are not in B. The following equalities hold:

$$\mathrm{A} = \{\mathrm{A} \cap \mathrm{B}, \mathrm{A} \cap \overline{\mathrm{B}}\}$$

$$\mathrm{B} = \{\mathrm{A} \cap \mathrm{B}, \overline{\mathrm{A}} \cap \mathrm{B}\}$$

$$\mathrm{A} \cup \mathrm{B} = \{\mathrm{A} \cap \mathrm{B}, \mathrm{A} \cap \overline{\mathrm{B}}, \overline{\mathrm{A}} \cap \mathrm{B}\}$$

A ∩ B is the set of species that are both in a and b, $\mathrm{A} \cap \overline{\mathrm{B}}$ the set of species that are in a but not in b, $\overline{\mathrm{A}} \cap \mathrm{B}$ the set of species that are in b but not in a, and A ∪ B contains all species that are either in a, in b, or in both a and b. The set A ∪ B is obtained by merging or mixing communities a and b altogether. Let $\mu$ be a real value in [0,1]. FAD increases by mixing if and only if the following inequality holds

$$\mathrm{FAD}(\mathrm{A} \cup \mathrm{B}) - \mu \times \mathrm{FAD}(\mathrm{A}) - (1-\mu) \times \mathrm{FAD}(\mathrm{B}) \geq 0$$

Knowing that

$$\mathrm{FAD}(\mathrm{A}) = \sum_{\mathrm{i,j} \in \mathrm{A} \cap \mathrm{B}} d_{ij} + \sum_{\mathrm{i,j} \in \mathrm{A} \cap \overline{\mathrm{B}}} d_{ij} + 2\sum_{\mathrm{i} \in \mathrm{A} \cap \mathrm{B}, \mathrm{j} \in \mathrm{A} \cap \overline{\mathrm{B}}} d_{ij}$$

$$\mathrm{FAD}(\mathrm{B}) = \sum_{\mathrm{i,j} \in \mathrm{A} \cap \mathrm{B}} d_{ij} + \sum_{\mathrm{i,j} \in \overline{\mathrm{A}} \cap \mathrm{B}} d_{ij} + 2\sum_{\mathrm{i} \in \mathrm{A} \cap \mathrm{B}, \mathrm{j} \in \overline{\mathrm{A}} \cap \mathrm{B}} d_{ij}$$

$$\mathrm{FAD}(\mathrm{A} \cup \mathrm{B}) = \sum_{\mathrm{i,j} \in \mathrm{A} \cap \mathrm{B}} d_{ij} + \sum_{\mathrm{i,j} \in \mathrm{A} \cap \overline{\mathrm{B}}} d_{ij} + \sum_{\mathrm{i,j} \in \overline{\mathrm{A}} \cap \mathrm{B}} d_{ij} + 2\sum_{\mathrm{i} \in \mathrm{A} \cap \mathrm{B}, \mathrm{j} \in \mathrm{A} \cap \overline{\mathrm{B}}} d_{ij} + 2\sum_{\mathrm{i} \in \mathrm{A} \cap \mathrm{B}, \mathrm{j} \in \overline{\mathrm{A}} \cap \mathrm{B}} d_{ij} + 2\sum_{\mathrm{i} \in \mathrm{A} \cap \overline{\mathrm{B}}, \mathrm{j} \in \overline{\mathrm{A}} \cap \mathrm{B}} d_{ij}$$

yields

$$\mathrm{FAD}(\mathrm{A} \cup \mathrm{B}) - \mu \times \mathrm{FAD}(\mathrm{A}) - (1-\mu) \times \mathrm{FAD}(\mathrm{B}) = (1-\mu)\sum_{\mathrm{i,j} \in \mathrm{A} \cap \overline{\mathrm{B}}} d_{ij} + \mu\sum_{\mathrm{i,j} \in \overline{\mathrm{A}} \cap \mathrm{B}} d_{ij} + 2(1-\mu)\sum_{\mathrm{i} \in \mathrm{A} \cap \mathrm{B}, \mathrm{j} \in \mathrm{A} \cap \overline{\mathrm{B}}} d_{ij} + 2\mu\sum_{\mathrm{i} \in \mathrm{A} \cap \mathrm{B}, \mathrm{j} \in \overline{\mathrm{A}} \cap \mathrm{B}} d_{ij} + 2\sum_{\mathrm{i} \in \mathrm{A} \cap \overline{\mathrm{B}}, \mathrm{j} \in \overline{\mathrm{A}} \cap \mathrm{B}} d_{ij} \geq 0$$

□

**${}^{q}\delta$ fulfills weak species monotonicity for all *q* in ]0, 1[ but not for *q* = 1, and species monotonicity for *q* = 0**

Let *N* be a set of *S*-1 species and *k* another species that is not included in *N*. Let **D** be the symmetric matrix of dissimilarities between all species in $N \cup \{k\}$. Consider **p** and **q** two vectors of length *S* containing the relative abundances of all species in $N \cup \{k\}$. In **q**, species *k* has zero abundance, while $p_i$ designates the relative abundance of any species *i* in *N*. In **p**, species *k* has a relative abundance equal to ε, and the relative abundance of any species $i \neq k$ is $(1-\varepsilon)p_i$. ${}^{q}\delta$ fulfills weak species monotonicity if and only if

$${}^{q}\delta(\mathbf{p}, \mathbf{D}) - {}^{q}\delta(\mathbf{q}, \mathbf{D}) \geq 0$$

For *q* = 0,

$${}^{0}\delta(\mathbf{p}, \mathbf{D}) - {}^{0}\delta(\mathbf{q}, \mathbf{D}) = 2\sum_{i \in N} d_{ik} \geq 0$$

The function fulfills species monotonicity, that is it increases with the addition of a species.

<u>For $q$ = 1,</u>

The condition for $^1\delta$ to increase with the addition of species $k$ is

$$^1\delta(\mathbf{p},\mathbf{D}) - {}^1\delta(\mathbf{q},\mathbf{D}) \geq 0$$

This condition corresponds to

$$\sum_{i\in N}\sum_{j\in N}(1-\varepsilon)^2 p_i p_j d_{ij} + 2\sum_{i\in N}(1-\varepsilon)p_i\varepsilon d_{ik} - \sum_{i\in N}\sum_{j\in N} p_i p_j d_{ij} \geq 0$$

$$2(1-\varepsilon)\varepsilon\sum_{i\in N} p_i d_{ik} \geq (1-(1-\varepsilon)^2)\sum_{i\in N}\sum_{j\in N} p_i p_j d_{ij}$$

$$2(1-\varepsilon)\varepsilon\sum_{i\in N} p_i d_{ik} \geq (1-1+2\varepsilon-\varepsilon^2)\sum_{i\in N}\sum_{j\in N} p_i p_j d_{ij}$$

$$2(1-\varepsilon)\varepsilon\sum_{i\in N} p_i d_{ik} \geq (2-\varepsilon)\varepsilon\sum_{i\in N}\sum_{j\in N} p_i p_j d_{ij}$$

$$\frac{2(1-\varepsilon)}{2-\varepsilon} \geq \frac{\sum_{i\in N}\sum_{j\in N} p_i p_j d_{ij}}{\sum_{i\in N} p_i d_{ik}}$$

Let $f(\varepsilon) = \frac{2(1-\varepsilon)}{2-\varepsilon}$, $f'(\varepsilon)=\frac{-2}{(2-\varepsilon)^2}<0$. $f$ is a decreasing function of $\varepsilon$. Knowing that $f(0) = 1$ and $f(1) = 0$, the existence of a value of $\varepsilon$ that fulfills the inequality $\frac{2(1-\varepsilon)}{2-\varepsilon} \geq \frac{\sum_{i\in N}\sum_{j\in N} p_i p_j d_{ij}}{\sum_{i\in N} p_i d_{ik}}$ depends of the value of $\frac{\sum_{i\in N}\sum_{j\in N} p_i p_j d_{ij}}{\sum_{i\in N} p_i d_{ik}}$. The function does not fulfill weak species monotonicity, that is there is no situation where the addition of a species will never decrease the diversity of a community. The increase or the decrease depends on the abundance at which it will be added and on its dissimilarities with present species. The inequality above shows that the possibility of diversity to increase with the addition of a new species implies that the new species has high effective originality ($\sum_{i\in N} p_i d_{ik}$ higher than $\sum_{i\in N}\sum_{j\in N} p_i p_j d_{ij}$). If instead the added species ($k$) is ordinary, sharing many similarities with species already present in a community, its addition, even at low abundance, may decrease the value of $^1\delta$.

<u>For $q$ in ]0,1[,</u>

$$
\begin{aligned}
{}^{q}\delta(\mathbf{p},\mathbf{D}) - {}^{q}\delta(\mathbf{q},\mathbf{D}) \\
&= \sum_{i\in N}\sum_{j\in N}(1-\varepsilon)^{2q} p_i^q p_j^q d_{ij} + 2\sum_{i\in N}(1-\varepsilon)^q \varepsilon^q p_i^q d_{ik} - \sum_{i\in N}\sum_{j\in N} p_i^q p_j^q d_{ij} \\
&= 2(1-\varepsilon)^q \varepsilon^q \left(\sum_{i\in N} p_i^q d_{ik}\right) - \left(1-(1-\varepsilon)^{2q}\right)\left(\sum_{i\in N}\sum_{j\in N} p_i^q p_j^q d_{ij}\right)
\end{aligned}
$$

As $2(1-\varepsilon)^q\varepsilon^q$, $\sum_{i\in N} p_i^q d_{ik}$, $(1-(1-\varepsilon)^{2q})$, and $\sum_{i\in N}\sum_{j\in N} p_i^q p_j^q d_{ij}$ are all nonnegative, the inequality to be verified

$$
2(1-\varepsilon)^q \varepsilon^q \left(\sum_{i\in N} p_i^q d_{ik}\right) - \left(1-(1-\varepsilon)^{2q}\right)\left(\sum_{i\in N}\sum_{j\in N} p_i^q p_j^q d_{ij}\right) \geq 0
$$

corresponds to

$$
\frac{(1-\varepsilon)^q \varepsilon^q}{(1-(1-\varepsilon)^{2q})} \geq \frac{1}{2}\frac{\sum_{i\in N}\sum_{j\in N} p_i^q p_j^q d_{ij}}{\sum_{i\in N} p_i^q d_{ik}}
$$

Let $f(\varepsilon) = \frac{(1-\varepsilon)^q\varepsilon^q}{(1-(1-\varepsilon)^{2q})}$,

$$
\begin{aligned}
f'(\varepsilon) &= \frac{(q(1-\varepsilon)^q\varepsilon^{q-1} - q(1-\varepsilon)^{q-1}\varepsilon^q)(1-(1-\varepsilon)^{2q}) - (1-\varepsilon)^q\varepsilon^q \times 2q(1-\varepsilon)^{2q-1}}{(1-(1-\varepsilon)^{2q})^2} \\
&= q(1-\varepsilon)^{q-1}\varepsilon^{q-1} \times \frac{(1-2\varepsilon)(1-(1-\varepsilon)^{2q}) - 2\varepsilon \times (1-\varepsilon)^{2q}}{(1-(1-\varepsilon)^{2q})^2} = \\
&= q(1-\varepsilon)^{q-1}\varepsilon^{q-1} \times \frac{1-2\varepsilon-(1-\varepsilon)^{2q}}{(1-(1-\varepsilon)^{2q})^2}
\end{aligned}
$$

The sign of $f'$ depends on $1-2\varepsilon-(1-\varepsilon)^{2q}$, with $q$ in $]0,1[$. Given that

$$
1-2\varepsilon-(1-\varepsilon)^{2q} = 1-2\varepsilon-(1-2\varepsilon+\varepsilon^2)^q
$$

$$
(1-2\varepsilon+\varepsilon^2)^q > 1-2\varepsilon+\varepsilon^2 > 1-2\varepsilon
$$

$f$ is thus a decreasing function of $\varepsilon$.

When $q$ = 0.5, $f(\varepsilon) = \frac{\sqrt{1-\varepsilon}}{\sqrt{\varepsilon}}$ and thus $f(\varepsilon) \xrightarrow[\varepsilon\to 0^+]{} \infty$. This means that there will always be a value of $\varepsilon$ sufficiently small so that $\frac{(1-\varepsilon)^q\varepsilon^q}{(1-(1-\varepsilon)^{2q})} \geq \frac{1}{2}\frac{\sum_{i\in N}\sum_{j\in N} p_i^q p_j^q d_{ij}}{\sum_{i\in N} p_i^q d_{ik}}$ with $q$ = 0.5.

For $q < 0.5$, as $1-(1-\varepsilon) \geq 1-(1-\varepsilon)^{2q}$, then $f(\varepsilon) \geq \frac{(1-\varepsilon)^q}{\varepsilon^{1-q}}$. Knowing that $\frac{(1-\varepsilon)^q}{\varepsilon^{1-q}} \xrightarrow[\varepsilon\to 0^+]{} \infty$, $f(\varepsilon) \xrightarrow[\varepsilon\to 0^+]{} \infty$. This means that there will always be a value of $\varepsilon$ sufficiently small so that $\frac{(1-\varepsilon)^q\varepsilon^q}{(1-(1-\varepsilon)^{2q})} \geq \frac{1}{2}\frac{\sum_{i\in N}\sum_{j\in N} p_i^q p_j^q d_{ij}}{\sum_{i\in N} p_i^q d_{ik}}$ with $0 < q < 0.5$.

For $q > 0.5$, let $g(\varepsilon) = (1-\varepsilon)^q\varepsilon^q$ and $h(\varepsilon) = (1-(1-\varepsilon)^{2q})$, so that $f(\varepsilon) = \frac{g(\varepsilon)}{h(\varepsilon)}$. We can use the Hospital rule to find the limit for $f(\varepsilon)$ when $\varepsilon \to 0^+$:

$$\lim_{\varepsilon\to 0^+} f(\varepsilon) = \lim_{\varepsilon\to 0^+} \frac{g'(\varepsilon)}{h'(\varepsilon)}$$

$$g'(\varepsilon) = -q(1-\varepsilon)^{q-1}\varepsilon^q + q(1-\varepsilon)^q\varepsilon^{q-1} = q(1-\varepsilon)^{q-1}\varepsilon^{q-1}(1-2\varepsilon)$$

$$h'(\varepsilon) = 2q(1-\varepsilon)^{2q-1}$$

$$\frac{g'(\varepsilon)}{h'(\varepsilon)} = \frac{(1-\varepsilon)^{-q}\varepsilon^{q-1}(1-2\varepsilon)}{2}$$

given that $q$ is in ]0.5, 1[, $q-1$ is negative which yields

$$\lim_{\varepsilon\to 0^+} f(\varepsilon) = \lim_{\varepsilon\to 0^+} \frac{g'(\varepsilon)}{h'(\varepsilon)} = \infty$$

This means that there will always be a value of $\varepsilon$ sufficiently small so that $\frac{(1-\varepsilon)^q\varepsilon^q}{(1-(1-\varepsilon)^{2q})} \geq \frac{1}{2}\frac{\sum_{i\in N}\sum_{j\in N} p_i^q p_j^q d_{ij}}{\sum_{i\in N} p_i^q d_{ik}}$ with $1 > q > 0.5$.

As a conclusion, the function ${}^q\delta$ thus fulfills weak species monotonicity for all $q$ in ]0,1[ and species monitonicity for $q = 0$.

□

**At the maximum of ${}^q\delta(\mathbf{p}, \mathbf{D})$ over p, a species has an effective originality equal to that of the other present species if *q* in ]0, 1[, and a species is either absent (has null abundance) or is present (has positive abundance) and has an effective originality equal to that of the other present species if *q* = 1.**

Recall that ${}^q\delta$ can be expressed as a mean of species effective originalities:

$$ {}^{q}\delta(\mathbf{p},\mathbf{D}) = \sum_{i=1}^{S} p_i \sum_{j=1}^{S} \frac{p_j^q}{p_i^{1-q}} d_{ij} = \sum_{i=1}^{S} p_i \, {}^{q}O_i(\mathbf{p},\mathbf{D}) $$

Any vector $\mathbf{p}_{max}$ that maximizes function ${}^{q}\delta(\mathbf{p},\mathbf{D})$ can be identified by resolving the Karush-Kuhn-Tucker (KKT) conditions

$$ \frac{\partial \, {}^{q}g(\mathbf{p},\mathbf{D})}{\partial p_i} = 0 $$

with ${}^{q}g(\mathbf{p},\mathbf{D}) = {}^{q}\delta(\mathbf{p},\mathbf{D}) - \lambda(\sum_{k=1}^{S} p_k - 1) + \sum_{k=1}^{S} \mu_k p_k$, $\lambda$ a constant satisfying $\lambda \neq 0$, and for all $k$, $\mu_k \geq 0$ and $\mu_k p_k = 0$. These conditions correspond to

$$ \frac{\partial \, {}^{q}\delta(\mathbf{p},\mathbf{D})}{\partial p_i} - \lambda + \mu_i = 0 $$

For weak monotone functions of diversity, for all $i$, $\mu_i = 0$. For all $i,j$ such that $\mu_i = \mu_j = 0$ and thus $p_i, p_j > 0$,

$$ \frac{\partial \, {}^{q}\delta(\mathbf{p},\mathbf{D})}{\partial p_i} = \lambda = \frac{\partial \, {}^{q}\delta(\mathbf{p},\mathbf{D})}{\partial p_j} $$

For all $p_i$,

$$ \frac{\partial \, {}^{q}\delta(\mathbf{p},\mathbf{D})}{\partial p_i} = \sum_{j=1}^{S} \frac{p_j^q}{p_i^{1-q}} d_{ij} + \sum_{k=1}^{S} p_k \frac{q \times p_i^{q-1}}{p_k^{1-q}} d_{ik} = \sum_{j=1}^{S} \frac{p_j^q}{p_i^{1-q}} d_{ij} + q \sum_{k=1}^{S} \frac{p_k^q}{p_i^{1-q}} d_{ik} $$
$$ = (q+1) \sum_{j=1}^{S} \frac{p_j^q}{p_i^{1-q}} d_{ij} = (q+1) \, {}^{q}O_i(\mathbf{p},\mathbf{D}) $$

Given that ${}^{q}\delta$ is weak species monotonous for all $q$ in ]0, 1[ but not for $q$ = 1, the set of all possible species is divided into two subsets at the maximum of ${}^{1}\delta(\mathbf{p},\mathbf{D})$: one where species' abundances render their effective originalities even, and the other where species have null abundances (knowing that the latter subset can be empty). For $q$ in ]0, 1[, at the maximum of ${}^{q}\delta(\mathbf{p},\mathbf{D})$, the abundances of all species in the set render their effective originalities even.

□

#### ${}^{0}\sigma(\mathbf{p},\mathbf{D})$ can decrease by the addition of a new species

Example: Let A be a set of three species with a constant dissimilarity, *d*, between them. Let B the set of these three species plus an additional fourth species distant from each of the former three by a dissimilarity $\delta$, with $\delta < d$. Consider *d* as the maximal possible dissimilarity: $d_{max} = d$.

$$ {}^{0}\sigma(\mathbf{p},\mathbf{D}) = \frac{\sum_{i=1}^{S}\sum_{j=1}^{S} d_{ij}}{S \times d_{max}} $$

${}^{0}\sigma$ applied to set A gives $\frac{3\times2\times d}{3\times d} = 2$

${}^{0}\sigma$ applied to set B gives $\frac{3\times2\times d+3\times2\times\delta}{4\times d} = \frac{3\times(d+\delta)}{2\times d}$

As soon as $\delta < \frac{d}{3}$, $\frac{3\times(d+\delta)}{2\times d} < \frac{3\times\frac{4d}{3}}{2\times d}$, which corresponds to $\frac{3\times(d+\delta)}{2\times d} < 2$.

□

#### ${}^{q}\sigma(\mathbf{p},\mathbf{D})$ is not weak species monotonous for *q* in ]0,0.5[

Let *N* be a set of *S*-1 species and *k* another species that is not included in *N*. Let **D** be the symmetric matrix of dissimilarities between all species in $N \cup \{k\}$. Consider **p** and **q** two vectors of length *S* containing the relative abundances of all species in $N \cup \{k\}$. In **q**, species *k* has zero abundance, while $p_i$ designates the relative abundance of any species i in *N*. In **p**, species *k* has a relative abundance equal to ε, and the relative abundance of any species $i \neq k$ is $(1-\varepsilon)p_i$. ${}^{q}\sigma$ fulfills weak species monotonicity if and only if

$$ {}^{q}\sigma(\mathbf{p},\mathbf{D}) - {}^{q}\sigma(\mathbf{q},\mathbf{D}) \geq 0 $$

$$\left( {}^{q}\sigma(\mathbf{p},\mathbf{D}) - {}^{q}\sigma(\mathbf{q},\mathbf{D}) \right) \times d_{max}$$
$$= S^{2q-1} \sum_{i \in N} \sum_{j \in N} (1-\varepsilon)^{2q} p_i^q p_j^q d_{ij} + 2S^{2q-1} \sum_{i \in N} (1-\varepsilon)^q \varepsilon^q p_i^q d_{ik}$$
$$- (S-1)^{2q-1} \sum_{i \in N} \sum_{j \in N} p_i^q p_j^q d_{ij}$$
$$= 2S^{2q-1}(1-\varepsilon)^q \varepsilon^q \left( \sum_{i \in N} p_i^q d_{ik} \right)$$
$$- \left( (S-1)^{2q-1} - S^{2q-1}(1-\varepsilon)^{2q} \right) \left( \sum_{i \in N} \sum_{j \in N} p_i^q p_j^q d_{ij} \right)$$

$$\left( {}^{q}\sigma(\mathbf{p},\mathbf{D}) - {}^{q}\sigma(\mathbf{q},\mathbf{D}) \right) \times d_{max} \xrightarrow[\varepsilon \to 0^+]{} -\left( (S-1)^{2q-1} - S^{2q-1} \right) \left( \sum_{i \in N} \sum_{j \in N} p_i^q p_j^q d_{ij} \right)$$

As $-\left((S-1)^{2q-1} - S^{2q-1}\right)\left(\sum_{i \in N} \sum_{j \in N} p_i^q p_j^q d_{ij}\right)$ is negative for $q$ in ]0,0.5[, ${}^{q}\sigma$ does not fulfill weak species monotonicity for all $q$ in ]0,0.5[.

□

**When 0 < *q* < 0.5, there exist some matrices D, such that the maximum of ${}^{q}\sigma(\mathbf{p},\mathbf{D})$ over p can be reached by a vector that contains at least one zero.**

Here is an example of such a matrix **D** (values below have been obtained with R package Rsolnp version 2.0.0, function csolnp, Galanos and Ye 2025, R Core Team 2025, and have been rounded with 4 digits):

Consider 3 species A, B, and C, with the following distances between them: $d_{AB} = 10$, $d_{BC} = d_{AC} = 2$.

<u>Below I consider as an example $q = 0.1$.</u> Let **D** be the matrix of distances between species A, B, and C and **p** = ($p_A$, $p_B$, $p_C$) the vector of relative abundances for the species. As ${}^{q}\delta$ is weak species monotonous and

$${}^{q}\sigma(\mathbf{p},\mathbf{D}) = \frac{{}^{q}\delta(\mathbf{p},\mathbf{D})}{S^{1-2q} \times d_{max}}$$

the maximum of ${}^{q}\sigma(\mathbf{D},\mathbf{p})$ over **p** is reached when species are split in two groups, "group 0" with species having zero abundance and "group +" with species having positive abundance that maximize ${}^{q}\delta(\mathbf{p}_{+}, \mathbf{D}_{+})$ over $\mathbf{p}_{+}$, with $\mathbf{p}_{+}$ the vector of relative abundance for species in group + and $\mathbf{D}_{+}$, the submatrix of **D** that contains only species in group +. While $S$ varies depending on the size of group +, in all cases $d_{max}$ is a constant equal to 10 in this case study.

With only 3 species, finding the maximum of ${}^{q}\sigma(\mathbf{p}, \mathbf{D})$ over **p** thus reduces to comparing the following two cases:

*Case 1. group + contains two species only and group 0 one species.*

{A, B} is the set of two species with the highest distance between them. Let $\mathbf{p}_{+}$ contain the relative abundance for species A and B and $\mathbf{D}_{+}$ be the submatrix of **D** that contains only species A and B. The maximum of ${}^{q}\delta(\mathbf{p}_{+}, \mathbf{D}_{+})$ over $\mathbf{p}_{+}$ is reached when $p_{\mathrm{A}} = p_{\mathrm{B}} = 0.5$.

$$^{0.1}\sigma\big((0.5,0.5,0), \mathbf{D}\big) = \frac{^{0.1}\delta\big((0.5,0.5,0), \mathbf{D}\big)}{2^{1-2q} \times 10} = 1$$

is thus the maximum value that ${}^{q}\sigma$ can reach, for **D** fixed, <u>over vectors of relative abundance that contain one zero and two positive values</u>.

*Case 2. group + contains all species A, B, C and group 0 is empty.*

The maximum of ${}^{q}\delta(\mathbf{p}, \mathbf{D})$ over **p** is reached when $p_{\mathrm{A}} = p_{\mathrm{B}} = 0.4345$ and $p_{\mathrm{C}} = 0.1310$.

$$^{0.1}\sigma\big((0.5,0,0.5,0), \mathbf{D}\big) = \frac{^{0.1}\delta\big((0.4345,0.4345,0.1310), \mathbf{D}\big)}{3^{1-2q} \times 10} = 0.9524$$

is thus the maximum value that ${}^{q}\sigma$ can reach, for **D** fixed, <u>over vectors of relative abundance that do not contain any zero</u>.

Thus, for our example, the maximum of ${}^{q}\sigma(\mathbf{D}, \mathbf{p})$ over **p** is reached when $p_{\mathrm{A}} = 0.5$, $p_{\mathrm{B}} = 0.5$, and $p_{\mathrm{C}} = 0$, that is when C is removed from the set and ${}^{q}\delta(\mathbf{p}, \mathbf{D})$ is maximized over **p** on the subset of species {A, B}.

#### ${}^{q}\boldsymbol{\sigma}(\mathbf{p}, \mathbf{D})$ is weak species monotonous for *q* in [0.5,1]

Let $N$ be a set of $S$-1 species and $k$ another species that is not included in $N$. Let **D** be the symmetric matrix of dissimilarities between all species in $N \cup \{k\}$. Consider **p** and **q** two vectors of length $S$ containing the relative abundances of all species in $N \cup \{k\}$. In **q**, species $k$ has zero abundance, while $p_i$ designates the relative abundance of any species $i$ in $N$. In **p**, species $k$ has a relative abundance equal to ε, and the relative abundance of any species $i \neq k$ is $(1-\varepsilon)p_i$. ${}^{q}\sigma$ fulfills weak species monotonicity if and only if

$$^{q}\sigma(\mathbf{p},\mathbf{D}) - {}^{q}\sigma(\mathbf{q},\mathbf{D}) \geq 0$$

For $q$ in ]0.5,1[, the validity of $^{q}\sigma(\mathbf{p},\mathbf{D}) - {}^{q}\sigma(\mathbf{q},\mathbf{D}) \geq 0$ comes directly from the fact that $^{q}\delta$ is weak species monotonous and $^{q}\sigma$ is an increasing function of species richness and $^{q}\delta$.

For $q = 0.5$, the validity of $^{q}\sigma(\mathbf{p},\mathbf{D}) - {}^{q}\sigma(\mathbf{q},\mathbf{D}) \geq 0$ comes from the fact that $^{q}\sigma = {}^{q}\delta/d_{max}$ with $d_{max}$ a constant, and $^{q}\delta$ is weak species monotonous.

**$^{q}H(\mathbf{p})$ is an index of species diversity than satisfies properties p0 to p3 for all $q$ in ]0, 1] and p4 for $q$ in ]0, 1[**

$$^{q}H(\mathbf{p}) = \sum_{i=1}^{S} \sum_{j=1, j\neq i}^{S} p_i^q p_j^q$$

p0. (nonnegativity). As $p_i \geq 0$ for all $i$, $^{q}H(\mathbf{p})$ also is $\geq 0$.

p1. (full concentration at minimum). If $p_k = 1$ and $p_j = 0$ for all $j \neq k$, then $^{q}H(\mathbf{p}) = 2\sum_{j=1, j\neq k}^{S} p_k^q p_j^q = 0$.

p4. (weak species monotonicity). True as soon as $^{q}\delta$ satisfies weak species monotonicity (for $q$ in ]0, 1[ see above). For $q = 1$, $^{1}H = 1 - \sum_{i=1}^{S} p_i^2$ is known as Gini-Simpson diversity (Gini 1912, Simpson 1949). Let $N$ be a set of $S$-1 species and $k$ another species that is not included in $N$. Consider **p** and **q** two vectors of length $S$ containing the relative abundances of all species in $N \cup \{k\}$. In **q**, species $k$ has zero abundance, while $p_i$ designates the relative abundance of any species $i$ in $N$. In **p**, species $k$ has a relative abundance equal to ε, and the relative abundance of any species $i \neq k$ is $(1-\varepsilon)p_i$. $^{1}H$ fulfills weak species monotonicity if and only if

$$^{1}H(\mathbf{p}) - {}^{1}H(\mathbf{q}) \geq 0$$

Knowing that

$$^{1}H(\mathbf{p}) - {}^{1}H(\mathbf{q}) = 1 - \sum_{i=1}^{S}(1-\varepsilon)^2 p_i^2 - \varepsilon^2 - 1 + \sum_{i=1}^{S} p_i^2$$

$$ {}^{1}H(\mathbf{p}) - {}^{1}H(\mathbf{q}) = (1 - 1 + 2\varepsilon - \varepsilon^2)\sum_{i=1}^{S} p_i^2 - \varepsilon^2 $$

$$ {}^{1}H(\mathbf{p}) - {}^{1}H(\mathbf{q}) = \varepsilon(2 - \varepsilon)\sum_{i=1}^{S} p_i^2 - \varepsilon^2 $$

$$ {}^{1}H(\mathbf{p}) - {}^{1}H(\mathbf{q}) = \varepsilon\left[(2 - \varepsilon)\sum_{i=1}^{S} p_i^2 - \varepsilon\right] $$

${}^{1}H(\mathbf{p}) \geq {}^{1}H(\mathbf{q})$ if and only if $\frac{2\sum_{i=1}^{S} p_i^2}{1+\sum_{i=1}^{S} p_i^2} \geq \varepsilon$. There always exists a value of $\varepsilon$ low enough to satisfy this inequality except when $\sum_{i=1}^{S} p_i^2 = 0$, that is when abundances are even in **q**: The addition of a species in a collection with evenly abundant species always decreases diversity as measured by Gini-Simpson index.

p2. (abundance evenness at maximum). Any vector $\mathbf{p}_{max}$ that maximizes function ${}^{q}H(\mathbf{p})$ can be identified by resolving the Karush-Kuhn-Tucker (KKT) condition

$$ \frac{\partial g(\mathbf{p})}{\partial p_i} = 0 $$

with $g(\mathbf{p}) = {}^{q}H(\mathbf{p}) - \lambda(\sum_{k=1}^{S} p_k - 1) + \sum_{k=1}^{S} \mu_k p_k$, $\lambda$ a constant satisfying $\lambda \neq 0$, and for all $k$, $\mu_k \geq 0$ and $\mu_k p_k = 0$.

$$ \frac{\partial\, {}^{q}H(\mathbf{p})}{\partial p_i} - \lambda + \mu_i = 0 $$

For weak monotone functions (see p4 above) of diversity, for all $k$, $p_k$ is positive and thus $\mu_k = 0$. This implies that for all $k,l$

$$ \frac{\partial\, {}^{q}H(\mathbf{p})}{\partial p_k} = \lambda = \frac{\partial\, {}^{q}H(\mathbf{p})}{\partial p_l} $$

$$ \frac{\partial \sum_{i=1}^{S} \sum_{j=1, j\neq i}^{S} p_i^q p_j^q}{\partial p_k} = 2q p_k^{q-1} \sum_{j=1, j\neq k}^{S} p_j^q $$

$\frac{\partial\, {}^{q}H(\mathbf{p})}{\partial p_k} = \frac{\partial\, {}^{q}H(\mathbf{p})}{\partial p_l}$ thus corresponds to

$$ p_k^{q-1} \sum_{j=1, j\neq k}^{S} p_j^q = p_l^{q-1} \sum_{j=1, j\neq l}^{S} p_j^q $$

$$\Leftrightarrow p_k^{q-1}\left(\sum_{j=1,j\neq k,l}^{S} p_j^q + p_l^q\right) = p_l^{q-1}\left(\sum_{j=1,j\neq k,l}^{S} p_j^q + p_k^q\right)$$

$$\Leftrightarrow p_l^{1-q} \sum_{j=1,j\neq k,l}^{S} p_j^q + p_l = p_k^{1-q} \sum_{j=1,j\neq k,l}^{S} p_j^q + p_k$$

$$\Leftrightarrow \left(\sum_{j=1,j\neq k,l}^{S} p_j^q\right)\left(p_l^{1-q} - p_k^{q-1}\right) + (p_l - p_k) = 0$$

$$\Leftrightarrow p_l = p_k$$

p3. (embedding of species richness). Given p2, $\max_{\mathbf{p}}\left({}^{q}H\right) = S^{1-2q}(S-1)$

$$\frac{\partial S^{1-2q}(S-1)}{\partial S} = (1-2q)S^{-2q}(S-1) + S^{1-2q} = S^{-2q}(2S - 2Sq - 1 + 2q)$$
$$= S^{-2q}(2(S-1)(1-q) + 1) > 0$$

$S^{1-2q}(S-1)$ is thus an increasing function of *S*. □

**${}^{q}h(\mathbf{p})$ is an index of species diversity than satisfies properties p0 to p4 for all *q* in ]0, 1]**

$${}^{q}h(\mathbf{p}) = S^{2q-1}\,{}^{q}H(\mathbf{p}) = S^{2q-1}\sum_{i=1}^{S}\sum_{j=1,j\neq i}^{S} p_i^q p_j^q$$

p0. (non-negativity). As $S^{2q-1} \geq 0$ and ${}^{q}H(\mathbf{p}) \geq 0$, ${}^{q}h(\mathbf{p})$ also is ≥ 0 for all **p**.

p1. (full concentration at minimum). If $p_k = 1$ and $p_j = 0$ for all $j \neq k$, then ${}^{q}H(\mathbf{p}) = 2\sum_{j=1,j\neq k}^{S} p_k^q p_j^q = 0$ and so is ${}^{q}h(\mathbf{p})$.

p2. (abundance evenness at maximum). Consider a subset of *Σ* species drawn from the *S* species of the collection. The maximum value of ${}^{q}H(\mathbf{p})$ in the subset of *Σ* species is obtained for $\mathbf{p}_{\Sigma\text{even}} = (1/\Sigma ... 1/\Sigma)^t$ and amounts to $\Sigma^{1-2q}(\Sigma - 1)$. Knowing that ${}^{q}h(\mathbf{p}_{\Sigma\text{even}}) = \Sigma - 1$ is an increasing function of *Σ* leads to ${}^{q}h$ maximized for $\mathbf{p}_{S\text{even}} = (1/S ... 1/S)^t$.

p3. (embedding of species richness) Given p2, $\max_{\mathbf{p}}\left({}^{q}h\right) = {}^{q}h(\mathbf{p}_{S\text{even}}) = S - 1$ is an increasing function of *S*. □

p4. (weak species monotonicity). True as soon as ${}^{q}\sigma$ satisfies weak species monotonicity, *i.e.*, for *q* in [0.5, 1] see above). ${}^{q}\sigma$ does not satisfy weak species monotonicity for *q* in ]0, 0.5[ see below

Let $N$ be a set of $S$-1 species and $k$ another species that is not included in $N$. Consider **p** and **q** two vectors of length $S$ containing the relative abundances of all species in $N \cup \{k\}$. In **q**, species $k$ has zero abundance, while $p_i$ designates the relative abundance of any species $i$ in $N$. In **p**, species $k$ has a relative abundance equal to ε, and the relative abundance of any species $i \neq k$ is $(1-\varepsilon)p_i$. ${}^{q}h$ fulfills weak species monotonicity if and only if

$${}^{q}h(\mathbf{p}) - {}^{q}h(\mathbf{q}) \geq 0$$

Knowing that

$$\begin{aligned} {}^{q}h(\mathbf{p}) - {}^{q}h(\mathbf{q}) \\ &= (1-\varepsilon)^2 S^{2q-1} \sum_{i=1}^{S} \sum_{j=1, j\neq i}^{S} p_i^q p_j^q + 2\varepsilon(1-\varepsilon) S^{2q-1} \sum_{i=1}^{S} p_i^q \\ &- (S-1)^{2q-1} \sum_{i=1}^{S} \sum_{j=1, j\neq i}^{S} p_i^q p_j^q \end{aligned}$$

$${}^{q}h(\mathbf{p}) - {}^{q}h(\mathbf{q}) = ((1-\varepsilon)^2 S^{2q-1} - (S-1)^{2q-1}) \sum_{i=1}^{S} \sum_{j=1, j\neq i}^{S} p_i^q p_j^q + 2\varepsilon(1-\varepsilon) S^{2q-1} \sum_{i=1}^{S} p_i^q$$

For $q$ in ]0, 0.5], $(1-\varepsilon)^2 S^{2q-1} - (S-1)^{2q-1}$ is negative. For $\varepsilon$ tending to zero, ${}^{q}h(\mathbf{p}) - {}^{q}h(\mathbf{q})$ thus takes negative values. □
